# Genomes from Bacteria Associated with the Canine Oral Cavity: a Test Case for Automated Genome-Based Taxonomic Assignment

**DOI:** 10.1101/577163

**Authors:** David A. Coil, Guillaume Jospin, Aaron E. Darling, Corrin Wallis, Ian J. Davis, Stephen Harris, Jonathan A. Eisen, Lucy J. Holcombe, Ciaran O’Flynn

## Abstract

Taxonomy for bacterial isolates is commonly assigned via sequence analysis. However, the most common sequence-based approaches (e.g. 16S rRNA gene-based phylogeny or whole genome comparisons) are still labor intensive and subjective to varying degrees. Here we present a set of 33 bacterial genomes, isolated from the canine oral cavity. Taxonomy of these isolates was first assigned by PCR amplification of the 16S rRNA gene, Sanger sequencing, and taxonomy assignment using BLAST. After genome sequencing, taxonomy was revisited through a manual process using a combination of average nucleotide identity (ANI), concatenated marker gene phylogenies, and 16S rRNA gene phylogenies. This taxonomy was then compared to the automated taxonomic assignment given by the recently proposed Genome Taxonomy Database (GTDB). We found the results of all three methods to be similar (25 out of the 33 had matching genera), but the GTDB approach was less subjective, and required far less labor. The primary differences in the remaining taxonomic assignments related to proposed taxonomy changes by the GTDB team.

## Introduction

With the ever-decreasing costs of DNA sequencing, it has become far easier/cheaper to sequence bacterial genomes than to analyze them. Understanding the gene content and metabolic pathways of a newly sequenced isolate is a time-consuming and knowledge-intensive process. Another, perhaps underappreciated, bottleneck is properly assigning taxonomy to a genome. This is most often seen with metagenome-assembled genomes (MAGs) which are unidentified prior to sequencing, but is even a problem with cultured isolates. Traditional morphological taxonomic assignment of bacterial isolates is tedious, and the more common approach of 16S rRNA gene PCR followed by Sanger sequencing is often uninformative beyond the genus level. Given the costs, some laboratories sequence isolate genomes directly, with no attempt at prior identification. While some have argued against the need for taxonomic assignment for many aspects of microbial genome analysis (e.g., [1][2]), there are many situations where taxonomy is considered valuable for making use of genomic information (e.g., see [3] and [4]).

There have been a number of proposed attempts to move to a genome-based taxonomy for bacteria and archaea, instead of relying on traditional chemotaxonomic/morphological characteristics of isolates [5][6][7]. These include the use of average nucleotide identity (ANI) [8][9][10][11], concatenated marker gene phylogenies (e.g. SILVA (unpublished) and GTDB (preprint: 10.1101/256800)) and shared protein content [12]. Most of these approaches however rely on a provisional identification (at least to genus), followed by locating/downloading the genomes of close relatives for comparison.

In this work we briefly describe the genome sequences of 33 bacterial isolates from the canine oral cavity. These isolates were collected as part of a larger project on canine oral health and had a preliminary taxonomy assigned through Sanger sequencing of the 16S rRNA gene [13][14]. After genome sequencing, we first assigned taxonomy to these isolates based on a manual combination of “whole genome” concatenated marker phylogenetic trees, average nucleotide identity (ANI), and 16S rRNA gene phylogenetic trees. We then compared this taxonomy to automated taxonomic assignments given by the recently proposed Genome Taxonomy Database (GTDB).

## Genomes and Taxonomy

### Genome selection

The study design, bacterial isolation, DNA extraction, isolation identification and genome sequencing/assembly have been previously described in our work on the *Porphyromonas* genus [15]. Briefly, bacterial isolates from the canine oral cavity were grown on supplemented Columbia Blood Agar containing 5% defibrinated horse blood (CBA; Oxoid, UK) with or without the addition of 5 mg/L Hemin (catalog no. H9039; Sigma) and 0.5 mg/L Menadione (catalog no. M5625; Sigma) or Heart Infusion Agar containing 5% defibrinated horse blood (HIA; Oxoid, UK) (Table 1). Aerobes were incubated at 38 °C under normal atmospheric conditions for 1-5 days. Microaerophilic and anaerobic strains were incubated at 38 °C for 1-21 days in a MACS1000 anaerobic workstation (Don Whitley, UK) with gas levels at 5% oxygen, 10% carbon dioxide, and 85% nitrogen for microaerophiles, and 10% hydrogen, 10% carbon dioxide, and 80% nitrogen for anaerobes. Following DNA extraction, library preparation, and Illumina sequencing, the reads were assembled using the A5-miseq assembly pipeline [16].

**Table 1:**
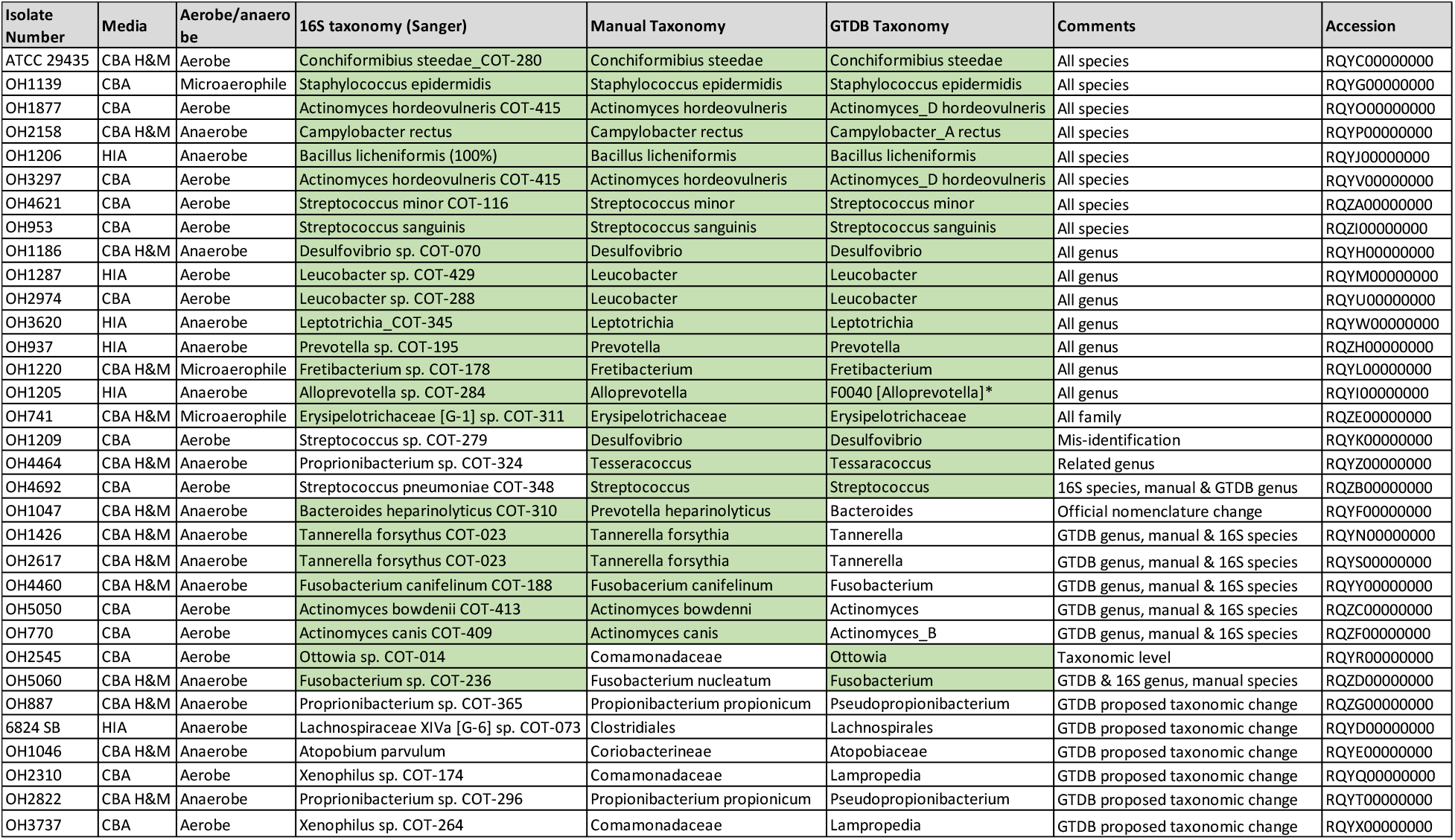
Comparative taxonomy of all 33 strains by three different methods. CBA: Columbia Blood Agar, HIA: Heart Infusion Agar, H&M: Hemin and Menadione. *“F0400” is actually a strain name, used a placeholder by GTDB when the authors believe that genome belongs in a new genus, for which no type representative is present.

The remaining *non-Porphyromonas* genomes were further screened by a combination of assembly metrics and CheckM [17] to estimate completeness and contamination. We chose for further study those that appeared to be the highest quality using admittedly somewhat arbitrary cutoffs of (a) fewer than 350 contigs in the assembly, (b) CheckM contamination score of <3%, and (c) CheckM completeness score of >90%. A subset of 33 genomes meeting these criteria was chosen to study in more detail.

### Preliminary taxonomic identification

All isolates were given a preliminary identification based on the Sanger sequencing of the 16S rRNA gene (F24/Y36 (9-29F/1525-1541R) primers). The 16S rRNA gene sequences were queried using BLAST [18] against the Canine Oral Microbiome Database (COMD). The COMD sequence database contained 460 published 16S rRNA gene sequences obtained from canine oral taxa (Genbank accession numbers <u>JN713151-JN713566</u> & <u>KF030193-KF030235</u> [13]. In addition, sequences were queried against the RDP 16S rRNA database v10_31 [19]. Species level identifications were made at 98.5%, genus at 94%, and family at 92%.

### Traditional taxonomic identification

The genomes were first uploaded to the RAST server [20] for annotation. The results archives were downloaded using the RAST API and then searched for full-length 16S rRNA gene sequences by scanning the annotations using the “SSU rRNA” tag and filtering for length greater than 1 kb. These sequences were uploaded to the Ribosomal Database Project (RDP) [19] and incorporated into their alignment. For first-pass taxonomic identification, an alignment was generated using SSU-ALIGN [21] of all ~12,000 type strain 16S rRNA gene sequences from RDP, along with our 33 sequences from RAST. This provided us with the general region of the tree for each isolate or group of isolates.

Next, we generated a series of concatenated “whole genome sequence” (WGS) marker trees for each isolate or group of isolates, at the taxonomic level determined by the 16S rRNA gene type strain tree. All available genome sequences from the genus or family of interest were downloaded from GenBank, except in cases where more than 500 genomes were available. In those genome-rich genera (*Bacillus, Clostridium, Streptococcus, Staphylococcus, Campylobacter*), the WGS trees were built with only a subset of genomes, selected from the type strain results. For example, *Staphylococcus* contains ~40 species, but only genomes from those species closest to our isolate of interest were needed to build a useful WGS tree.

For all WGS trees, an outgroup genome(s) was chosen from nearby in the NCBI taxonomy (e.g. another genus in the same family). The file names and sequences were reformatted for easier visualization. The assemblies were then screened for 37 core marker genes [22] using PhyloSift [23] in the search and align mode using “isolate” and “besthit” flags. PhyloSift concatenates and aligns the hits of interest, then the sequences are subsequently extracted from the PhyloSift output files and added to a single file for tree-building. An approximately maximum-likelihood tree was then inferred using FastTree2 with default parameters [24].

For all isolates where the WGS tree indicated a possible placement into a well-defined clade with sequenced genomes, we downloaded the type strain genome sequences for every member of that genus/family from Refseq at NCBI. These were used to create an ANI (average nucleotide identity) matrix using FastANI (preprint: 10.1101/225342) for each group and anything having an ANI >95% to a type strain was considered to belong to that species [8][25].

For the majority of taxa, the WGS tree and ANI matrix were still inadequate to assign taxonomy, due to a paucity of genome sequences for many groups. In those cases, we were forced to rely solely on the 16S rRNA gene for taxonomy. This was again accomplished through RDP, by downloading all sequences for a given genus/family and looking for well-supported placement into a monophyletic clade. For all groups with fewer than 3000 sequences, the alignment was downloaded directly from RDP, for larger groups the sequences were downloaded and the alignment generated with SSU-ALIGN. All alignments were cleaned to remove problematic characters in the headers using a custom script [26] and all trees were inferred using FastTree2 with default parameters.

Final taxonomic assignments were based on a taxa-dependent combination of the WGS trees, the ANI results, and the 16S rRNA gene trees (Table 1). First priority was given to the ANI results, then to placement within a well-supported WGS tree with monophyletic clades, and then finally to 16S rRNA gene-based results. As a result of inadequate mapping between phylogeny and taxonomy, some isolates were only assigned to the genus or family level, and in one case the order level. Examples of both informative and non-informative WGS/16S trees, along with a sample ANI matrix, can be found in the Supplemental Materials.

### Genome Taxonomy Database

The Genome Taxonomy Database (GTDB) is a recently proposed system attempting to create a genomic-based, standardized bacterial taxonomy (preprint: 10.1101/256800). The creators chose 120 single-copy conserved marker genes to generate a tree of 94,759 genomes, and used the topology of this tree to propose large-scale revisions to bacterial taxonomy. During our work on this project, the authors released a tool called “GTDB-Tk” [27] which attempts to automate taxonomic assignment for genomes, based on their revised GTDB taxonomy. The tool uses a combination of phylogenetic tree topology, ANI values, and relative evolutionary divergence (RED). We screened our 33 isolates using this tool and compared the taxonomic assignments to the manual process above (Table 1). Note that an alphabetic suffix in the GTDB taxonomy (e.g. “*Actinomyces_B*”) indicates that, within the GTDB tree, the genus does not belong to a monophyletic group with the type species of that genus.

## Discussion

Here we present the genomes of 33 bacterial isolates from the canine oral cavity. Some are from groups known to be involved in human and canine oral health (e.g. *Actinomyces* and *Fusobacterium*) and others have not been previously suggested to play such a role. A few of our isolates appear to be novel and potentially represent new species or genera within their groups. For example Canine Oral Taxon number 073 (COT073) and bacterial isolate number OH741 were only identified to the order and family level respectively.

The difference in labor required by the two genomic methods of taxonomic assignment was noticeable, with the manual method having taken two weeks of daily downloads, alignments, ANI comparisons, and tree-building, whereas the GTDB automated method ran overnight. The latter also is much less subjective (for the user) than the former. A comparison of the three approaches to taxonomic assignments shows a very high degree of similarity (Table 1).

For 48.5% (16/33) of isolates the three methods gave identical results; for 18.2% of isolates the 16S and manual methods gave the same results; for 9.1% of isolates the manual and GTDB methods gave the same results; and for 6.1% of isolates the 16S and GTDB methods gave the same results. Finally, for 18.2% of isolates all three methods gave different results to each other i.e. different genera within same family or different levels of assignment (granularity/resolution) within the same branch of the tree. Most of the differences between the GTDB taxonomy and the other approaches was due to proposed taxonomic revisions by the GTDB group. For example COT073 was placed within the Clostridiales order in the manual taxonomy, but that order was subdivided into several new groups in the GTDB taxonomy, based on calculations of relative evolutionary divergence (RED) within that group.

Our results suggest that the use of the automated GTDB tool to assign taxonomy to unidentified bacterial isolates is less subjective and much faster than manual assignment, while giving very similar results (e.g. identical taxonomy, difference only in taxonomic rank, or proposed taxonomic changes). Both genomic methods offer improved taxonomic resolution relative to 16S Sanger sequencing, at the additional cost/burden of requiring the entire genome. Finally, we expect these genomes will be of use to researchers studying canine oral health since the vast majority of closely related isolates so far sequenced have been collected from human hosts.

## Accession numbers

The whole-genome shotgun projects have been deposited at DDBJ/ENA/GenBank under the accession numbers RQYC00000000-RQZI00000000. The versions described in this paper are versions RQYC01000000-RQZI01000000. These strains were submitted to NCBI using our “manual taxonomy” assignments, since the GTDB taxonomy is not yet widely accepted. However, for some of the isolates, NCBI made their own minor taxonomic revisions.

## Acknowledgements

The authors would like to acknowledge the laboratory staff at WALTHAM Centre for Pet Nutrition for growing the bacterial isolates, extracting genomic DNA, and for initial species identification. The authors would also like to acknowledge John Zhang from UC Davis for the sequencing library preparation of isolates. The sequencing was carried by the DNA Technologies and Expression Analysis Cores at the UC Davis Genome Center, supported by NIH Shared Instrumentation Grant 1S10OD010786-01. This work was funded by WALTHAM Centre for Pet Nutrition.

---

**Supplemental Figure 1:**
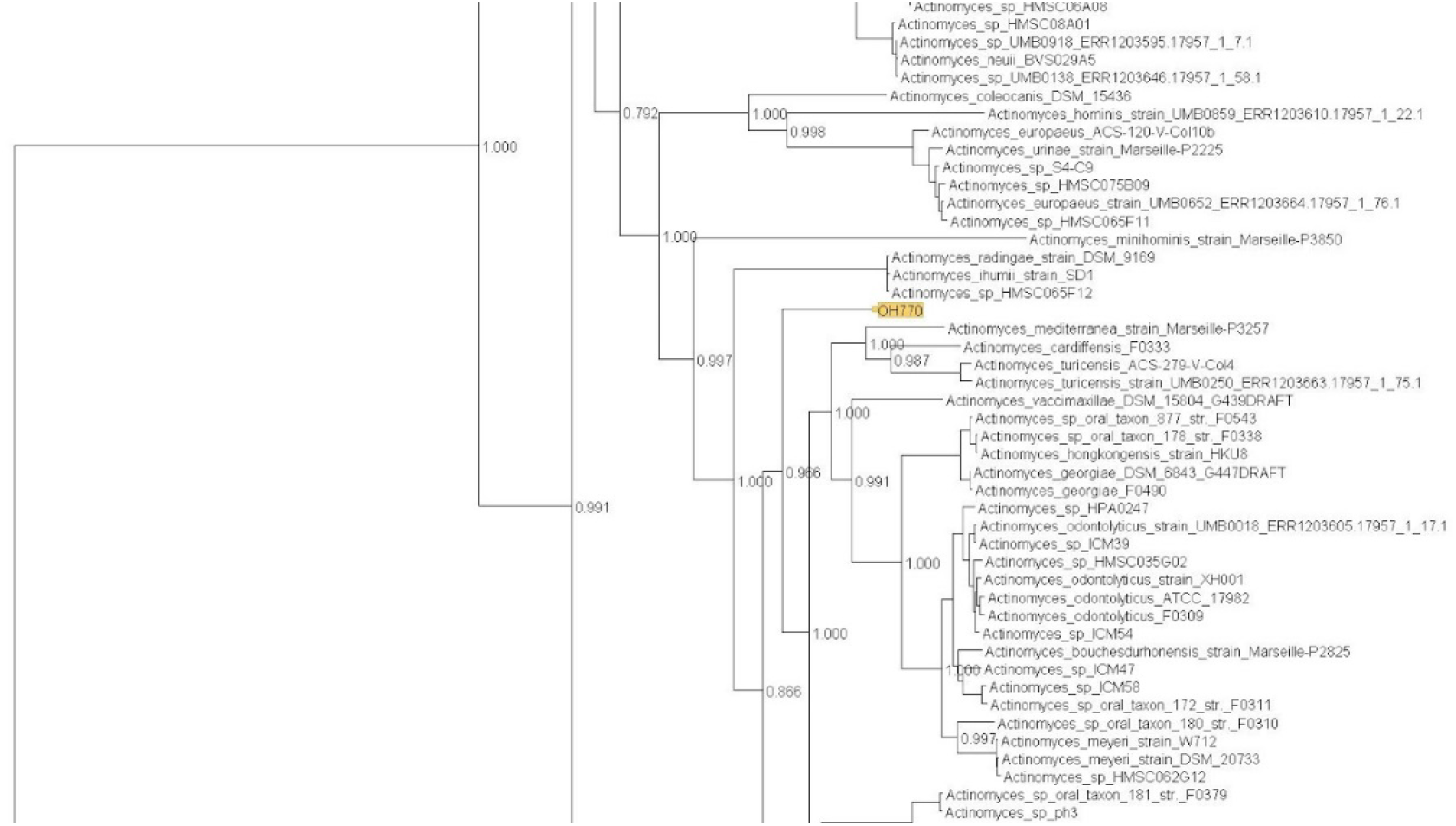
A taxonomically inconclusive WGS tree for isolate OH770. The isolate is not found within a clade of other sequenced isolates with species-level taxonomy.

**Supplemental Figure 2:**
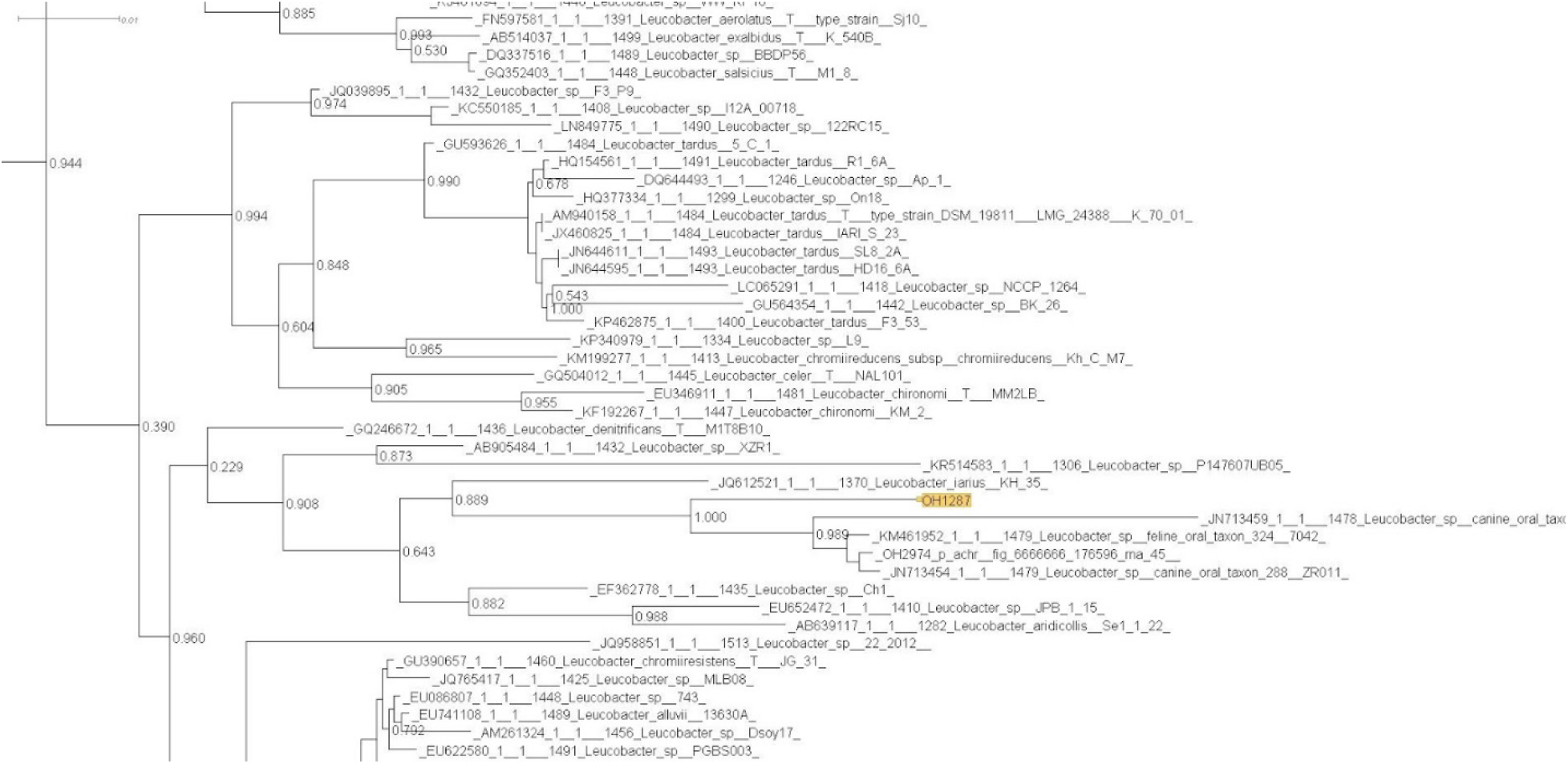
A taxonomically inconclusive 16S rRNA gene tree for isolate OH1287. Taxonomy is not congruent with phylogeny and many neighboring sequences are only identified to the genus level.

**Supplemental Figure 3:**
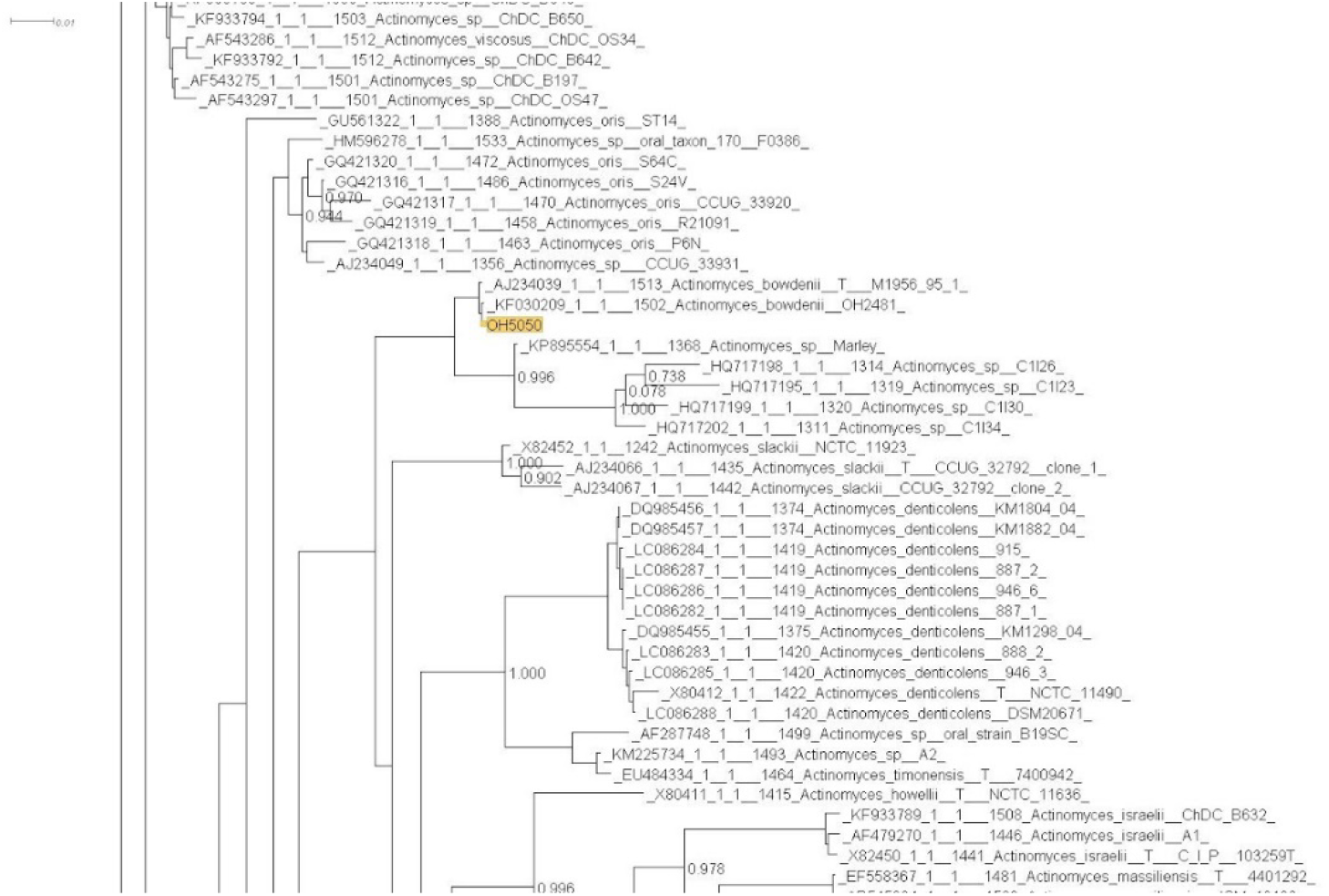
A taxonomically informative 16S rRNA gene tree for isolate OH5050. The isolate is found in a monophyletic clade and the name given to the closest relatives is not found elsewhere in the tree.

**Supplemental Table 1:**
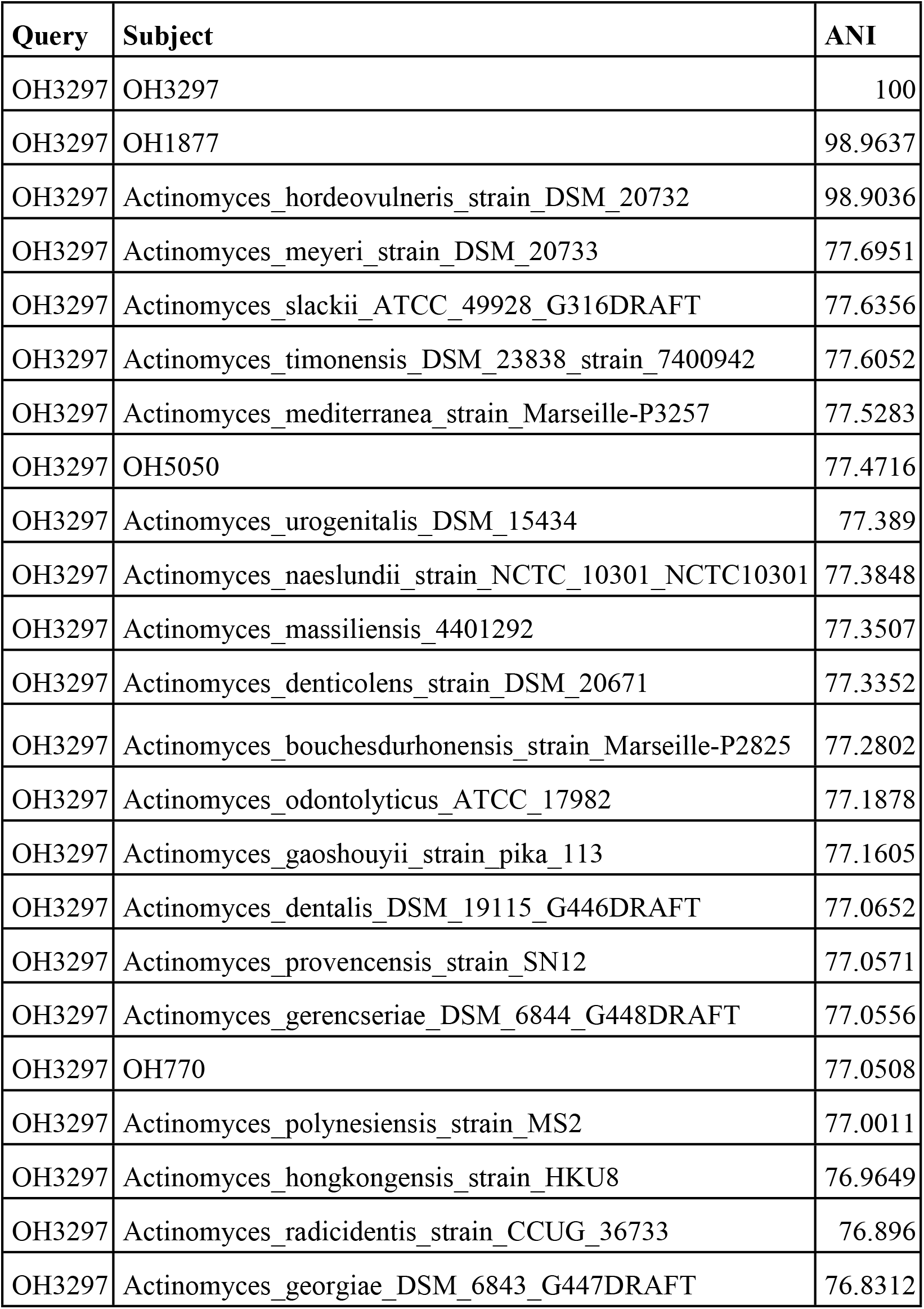
A sample ANI result for isolate OH3297. The isolate is ~99% identical to both OH1877 and to *Actinomyces hordeovulneris*. The ANI drops off quite rapidly to other members of the genus.

